# Familiarity Facilitates Feature-based Face Processing

**DOI:** 10.1101/058537

**Authors:** Matteo Visconti di Oleggio Castello, Kelsey G. Wheeler, Carlo Cipolli, M. Ida Gobbini

**Author notes:** Correspondence: M. Ida Gobbini,;, Matteo Visconti di Oleggio Castello.

## Abstract

Recognition of personally familiar faces is remarkably efficient, effortless and robust. We asked if feature-based face processing facilitates detection of familiar faces by testing the effect of face inversion on a visual search task for familiar and unfamiliar faces. Because face inversion disrupts configural and holistic face processing, we hypothesized that inversion would diminish the familiarity advantage to the extent that it is mediated by such processing. Subjects detected personally familiar and stranger target faces in arrays of two, four, or six face images. Subjects showed significant facilitation of personally familiar face detection for both upright and inverted faces. The effect of familiarity on target absent trials, which involved only rejection of unfamiliar face distractors, suggests that familiarity facilitates rejection of unfamiliar distractors as well as detection of familiar targets. The preserved familiarity effect for inverted faces suggests that facilitation of face detection afforded by familiarity reflects mostly feature-based processes.

## Introduction

Humans are thought to be face-experts. We are able to draw important information from faces such as emotions from facial expressions [1,2], direction of attention from eye gaze and head position [1,2], and recognition of identity [3], [4]. When focusing on face identity, human performance is dramatically different for familiar and unfamiliar faces. Despite the subjective impression of efficient or “expert” perception of faces in general, performance accuracy when discriminating unfamiliar face identities or perceiving that different images are of the same unfamiliar identity are markedly worse than for familiar faces [3,5–12].

In previous work, we showed that personally familiar faces have a more robust representation as compared to unfamiliar faces for both early detection and perception of social cues. Familiar as compared to unfamiliar faces can be detected with reduced attentional resources and can be processed without conscious awareness [13]. Moreover, social cues, such as eye-gaze or head orientation, are processed faster when conveyed by familiar faces [14]. With a saccadic reaction paradigm, we found that participants were able to detect and shift their gaze to familiar faces in 180 ms [15] when the distractors were faces of strangers, a latency shorter than the known evoked potentials that differentiate familiar from stranger faces [16]; but see [17]). Overall, these results highlight a difference in processing between familiar and unfamiliar faces and point to a facilitation of familiar face processing that precedes the activation of a conscious, view-invariant representation [13,15], and that extends to the local features of a familiar face [12,14].

In order to test the hypothesis that fast and efficient detection of familiar faces relies primarily on feature-based processing, we assessed whether the advantage for familiar face detection persists for inverted faces. Face inversion has been used to demonstrate face-specific processing and the role of configural processing when faces are presented upright. Inverting a face disrupts configural and holistic processing, thereby increasing reliance on parts-based processing [18–31]). Faces are characterized by two types of relational/configurational properties: first-order relational properties (e.g.; eyes above the nose above the mouth) and second-order relational properties (e.g. spacing between the eyes) [32–34]. Another term used in the face literature for face processing is “holistic” [35], meaning that all face-parts are processed as a whole [36]. In the present experiment we hypothesized that if familiar face recognition exploits identity-specific local facial features, then the advantage for personally familiar faces should be maintained with face inversion. On the other hand, if familiar face recognition relies on holistic or configural processing, face inversion should eliminate the familiarity advantage. We used a visual search task for personally familiar and unfamiliar identities with upright and inverted faces. The results showed that the advantage for familiar faces persists also after inversion. We discuss these findings in terms of parts-based processing for efficient detection of personally familiar faces.

## Methods

Raw data, analysis scripts, and presentation code are available at [LINK OMITTED WHILE UNDER REVIEW]

### Participants

19 subjects (12 male, mean age: 24.79, SD 3.71) from three groups of friends participated in the experiment. No formal power estimate was computed to determine sample size, but we aimed for a sample size that was larger than that in a paper by Tong & Nakayama (1999, 8-16 subjects) on a visual search task for one’s own face, while recruiting subjects that were highly familiar with the familiar stimuli. We chose friends that had extensive daily interaction with each other occurring for at least one year prior to the experiment. They were recruited from the Dartmouth College graduate and undergraduate community. All had normal or corrected-to-normal vision. Subjects were reimbursed for their participation; all gave written informed consent to use their pictures for research and to participate in the experiment in accordance with the Declaration of Helsinki. The Dartmouth Committee for the Protection of Human Subjects approved the experiment (Protocol 21200).

### Stimuli

For each subject we created three sets of images: target familiar faces (two identities: one male, one female), target stranger faces (two identities: one male, one female), and distractor stranger faces (twelve identities: 6 male, 6 female). Prior to the experiment, subjects and their friends had their pictures taken to be used as stimuli in the experiment. To ensure that all stimuli were of equal image quality, pictures were taken in a photo studio with standardized lighting, camera placement and camera settings. For each identity we used two different pictures taken in the same session to reduce image-specific learning. The familiar targets were chosen among the subject’s friends. The pictures of the 14 stranger individuals (12 distractor identities and 2 target identities) were taken at the University of Vermont with the same lighting, camera placement and settings as used for subjects recruited at Dartmouth College. For each subject the two unfamiliar target identities were chosen randomly. Inverted stimuli were created by rotating the images 180°. Images were cropped and converted to grayscale using custom code written in Python on Mac OS X 10.9.5. The average pixel intensity of each image (ranging from 0 to 255) was set to 128 with a standard deviation of 40 using the SHINE toolbox (function *lumMatch*) [37] in MATLAB (R2014a).

Stimuli for visual search trials consisted of two, four, or six face images positioned on the vertices of a regular hexagon centered on the fixation point, such that the center of each image was 7° of visual angle from the fixation point. Each image subtended 4° x 4° of visual angle. The position of the stimuli always created a shape symmetrical with respect to the fixation point (see Fig 1). All face images for each block were either upright or inverted.

**Fig 1.**
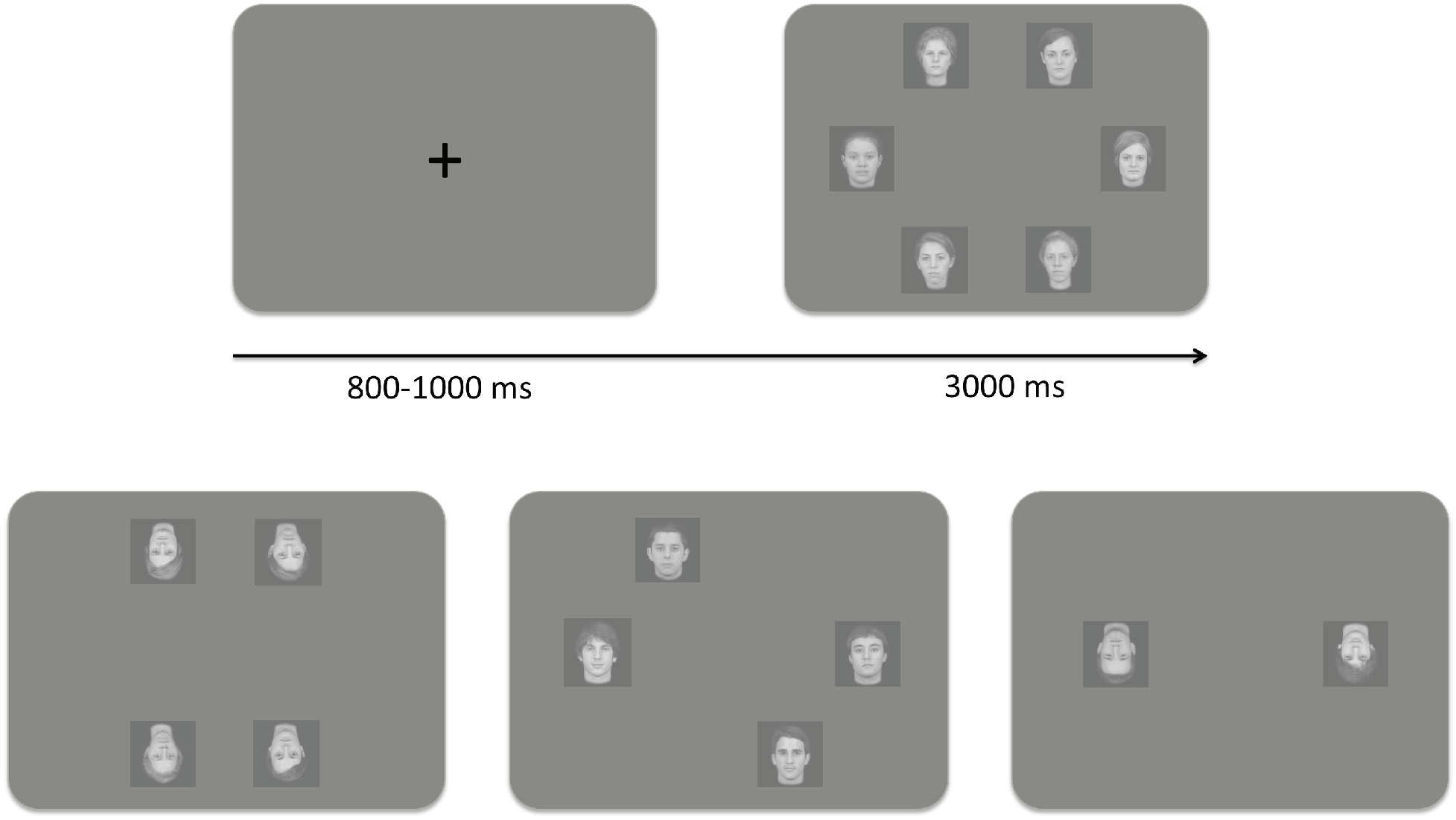
Experimental paradigm and example of the stimuli. On each trial, a central fixation cross appeared for a jittered period between 800-1000 ms, followed by a visual search array of two, four, or six faces displayed for a maximum of three seconds.

### Experimental setup

The experiment was run on a GNU/Linux workstation (Xubuntu 14.04 with low-latency kernel 3.13, CPU AMD FX-4350 quad-core 4.2 GHz, 8GB RAM, AMD Radeon R9 270 video card with radeon drivers) and a DELL 2000FP screen, set at a resolution of 1600×1200 pixels with a 60hz refresh rate, using Psychtoolbox (version 3.0.12) in MATLAB (R2014b). Subjects sat at a distance of approximately 50 cm from the screen (eyes to screen) in a dimly lit room.

### Task

Subjects were briefly familiarized with the images used in the visual search task before starting the experiment. Images (both upright and inverted) were presented in random order. Each image was presented for two seconds. After the image disappeared, subjects were required to press a key to continue to the next image. They were instructed to carefully observe each face for the entire presentation and to continue at their own pace.

The visual search session consisted of eight blocks, with a short break after the first four blocks. In each block, subjects were instructed to search for one of the four target identities, with one upright and one inverted block for each identity. Within each block, all distractor faces were of the same sex and in the same orientation as the target images. Subjects responded as quickly and accurately as possible by pressing either the left-arrow key (target present) or the right arrow-key (target absent). They received feedback (a beep) if they responded incorrectly or did not respond within three seconds. No feedback was given for correct answers. Eye movements were explicitly allowed.

The order of blocks was counterbalanced for familiarity and face orientation within each subject. Familiarity always changed from one block to the next, while inversion changed every two blocks. Because of software error, the sex of the targets wasn’t counterbalanced across subjects: 12/19 subjects had male targets in the first half of the experiment and female targets in the second half (and the converse for the remaining 7/19 subjects).

Each block started with 24 practice trials followed immediately by 120 test trials. At the beginning of each block, subjects were shown the target identity (upright or inverted) and pressed a key to start the block. On each trial, a central fixation cross appeared for a jittered period between 800-1000 ms, followed by a visual search array of two, four, or six faces displayed for a maximum of three seconds.

Target images appeared in half of the trials. The target was equally likely to appear in the left or right hemifield to avoid possible lateralization biases. Distractor faces were randomly chosen from the set of six distractor identities, and all distractors were different from each other.

Stranger target identities never appeared as distractors. Each trial type was repeated 10 times in each block (with distractors randomly sampled every time). Each block thus had 120 trials: 3 (Set Size) × 2 (Target Presence) × 20 (2 different target images × 10 repetitions). The order of the trials within each block was randomized.

### Statistical Analyses

The analyses were run in R (version 3.2.3). The code for all the analyses are available on the Open Science Framework website ([LINK OMITTED WHILE UNDER REVIEW]) as RMarkdown notebooks.

To assess statistical significance we fitted Generalized Mixed Models using the package *lme4* (version 1.1.11 [38]). Significance of the model parameters was tested using a Type 3 analysis of deviance (Wald’s *χ*^2^ test), as implemented in the package *car* ([39], version 2.1.1). We also used the following additional packages in our analyses:

- *dplyr* (version 0.4.3, [40]
- *ggplot2* (version 2.1.0, [41]
- *foreach* (version 1.4.3, [42]
- *doParallel* (version 1.0.10, Analytics and Weston, 2015b)
- *knitr* (version 1.12.3, [43–45]
- *assertthat* (version 0.1, Wickham, 2013)
- *broom* (version 0.4.0, Robinson, 2015)

We analyzed subjects’ accuracies using Logit Mixed Models [46], and reaction times of correct trials only with Linear Mixed Models, separately for target present and target absent trials. For each model, we entered Set Size, Familiarity, and Target Orientation as main effects with all their interactions. The initial random-effect structure contained both subjects and items terms.

For the latter term we entered the combination of stimuli appearing on the screen regardless of their position. This allowed us to model the variance due to subject and item (specific images) differences. We also added an extra regressor that indicated the sex of the target, and added random slopes with respect to this term for both subjects and items. We considered this term as a covariate, and thus we didn’t analyze it further.

The initial random-effect structure was tested using a log-likelihood ratio test against reduced models (created by removing random slopes first). For the linear models on reaction times in both target present and absent trials, the final structure contained subjects with random slopes and intercepts, and items with random intercepts—the model with random slopes for items failed to converge, thus we used a less complex model. The final logit models on accuracies in target present trials had subjects with random intercepts only, while in target absent trials it had subjects with both random intercepts and slopes.

After fitting the models with zero-sum contrasts for the regressors, we tested statistical significance of the fixed-effect terms using a Type 3 analysis of deviance (Wald’s *χ*^2^ test), as implemented in the package *car* [39]. For the models on reaction times we log-transformed the independent variable to account for the skewness of the distribution of reaction times; visually inspecting the predicted vs. residual plot confirmed that such a transformation provided a better fit for the model. The final linear model was refitted using restricted maximum likelihood estimation (REML).

We used a bootstrapping procedure [47] to investigate the direction of the significant effects found by the models. Trials were always bootstrapped maintaining the structure of the original dataset. For example, for any bootstrap sample the number of trials within each subject and condition (Set Size, Target Presence, Target Orientation, Familiarity, and Target Sex) was preserved, and trials were sampled with replacement only within the appropriate subject and condition. For the next sections, numbers in square brackets represent 95% basic bootstrapped confidence intervals (CI) after 10,000 replications.

We also estimated Set Size 1 intercept and search slopes—which provide information about target-recognition and distractor-rejection processes (Tong & Nakayama 1999)—by fitting a regression line for each subject and condition separately. To obtain 95% confidence intervals we bootstrapped the trials (in a stratified fashion, i.e., maintaining the factorial design of the conditions) and ran the regression model again, repeating this process 10,000 times.

## Results

### Accuracy

#### Target Present Trials

Subject responses were overall highly accurate, with average accuracy in target present trials of *93.29%* CI: [92.80, 93.78] (see Fig 2). We found a significant main effect of set size (χ^2^ (2) = 75.01, p < .001) and of target orientation (*χ*^2^ (1) = 19.37, p < .001). Subjects were more accurate when fewer distractors appeared on the screen (one distractor 96.09% [95.43, 96.74]; three distractors 93.62% [92.76, 94.41]; and five distractors 90.16% [89.14, 91.15]), and when faces were presented upright (upright 94.69% [94.06, 95.31]; inverted 91.89% [91.12, 92.63]). S1 File shows the *χ*^2^ values for the other main and interaction terms.

**Fig 2.**
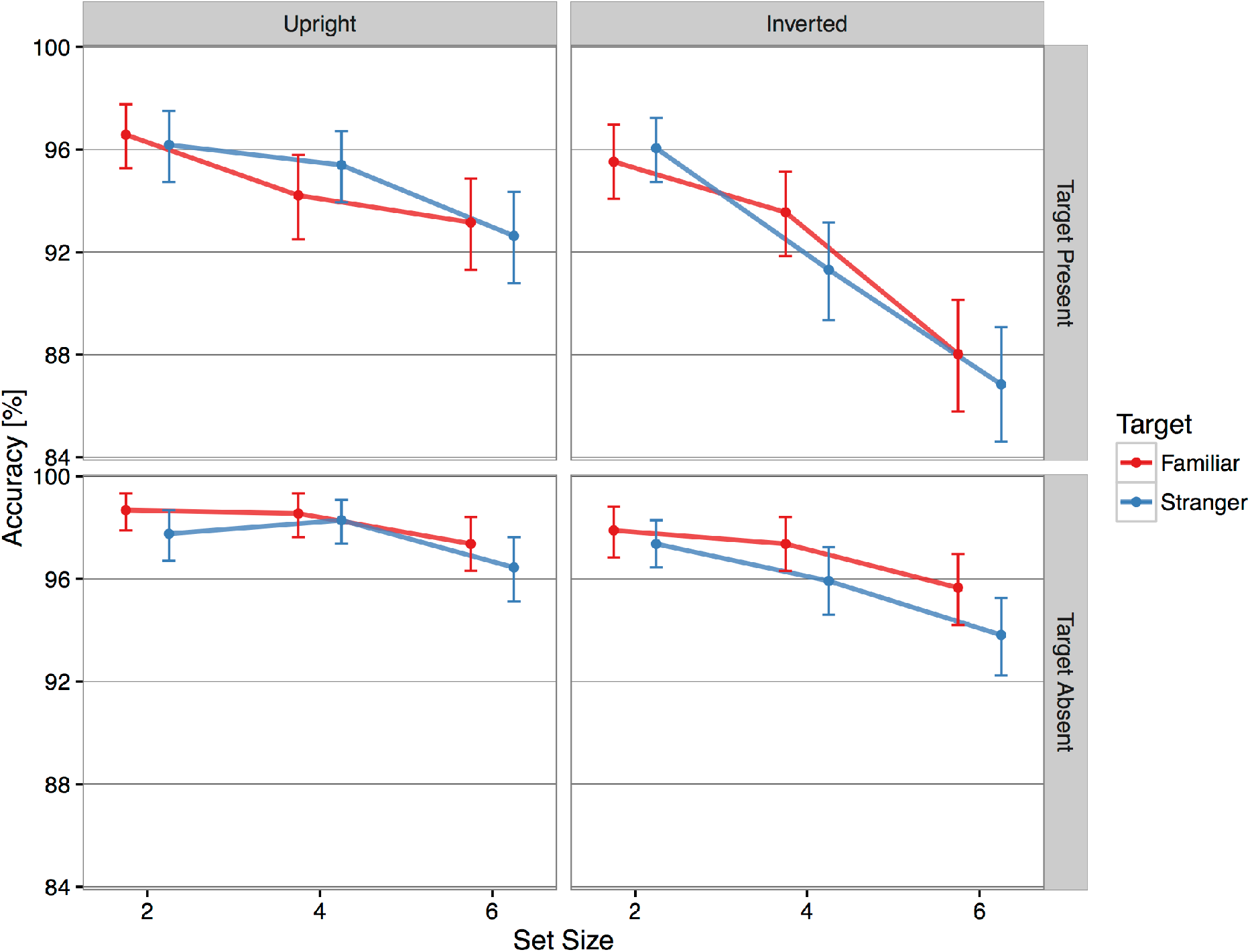
Average accuracy according to target orientation (columns) and presence of the target (rows). Subjects were overall highly accurate, with better performance when faces were presented upright and with fewer distractors. Red: familiar targets; Blue: stranger targets. Error bars show 95% bootstrapped confidence intervals.

#### Target Absent Trials

Subject responses were also highly accurate on target absent trials, with average accuracy of 97.09% [96.78, 97.41]. We found a significant main effect of set size (*χ*^2^ (2) = 25.54, p < .001), familiarity (*χ*^2^ (1) = 6.75, p < .01), and target orientation (*χ*^2^ (1) = 16.54, p < .001) but no other significant main or interaction effects (see S1 File). Subjects were more accurate at saying the target was absent when looking for a familiar face (familiar 97.59% [97.15, 98.00]; stranger 96.60% [96.10, 97.08]) and when faces were presented upright (upright 97.85% [97.43, 98.25]; inverted 96.34% [95.83, 96.82]). Subjects’ accuracy was lower with six distractors (two distractors 97.93% [97.47, 98.39]; four distractors 97.53% [97.01, 98.03]; and six distractors 95.82% [95.16, 96.48]).

### Reaction Times

#### Target Present Trials

All main effects of interest were statistically significant: Set Size (*χ*^2^ (2) = 1318.93, p < .001), Familiarity (*χ*^2^ (1) = 169.61, p < .001), and Target Orientation (*χ*^2^ (1) = 400.49, p < .001). We found significant interactions of Set Size x Familiarity (*χ*^2^ (2) = 8.59, p < .05) reflecting faster reaction times for familiar face trials; of Familiarity x Target Orientation (*χ*^2^(1) = 9.16, p < .001) reflecting a larger familiarity effect for upright faces, and Set Size x Familiarity x Target Orientation (*χ*^2^(2) = 11.17, p < .001) reflecting mostly a difference in the effect of familiarity on slopes for upright versus inverted faces (see S1 File).

Subjects were overall faster when searching for a familiar face than a stranger face, and they were faster with upright faces than inverted faces (see Fig 3). The advantage for familiar faces was 114 ms [97, 131] in the upright condition, and 75 ms [55, 95] in the inverted condition, with a difference of 39 ms [13, 65]). Fig 4 shows the effect size of Familiarity at each set size.

**Fig 3.**
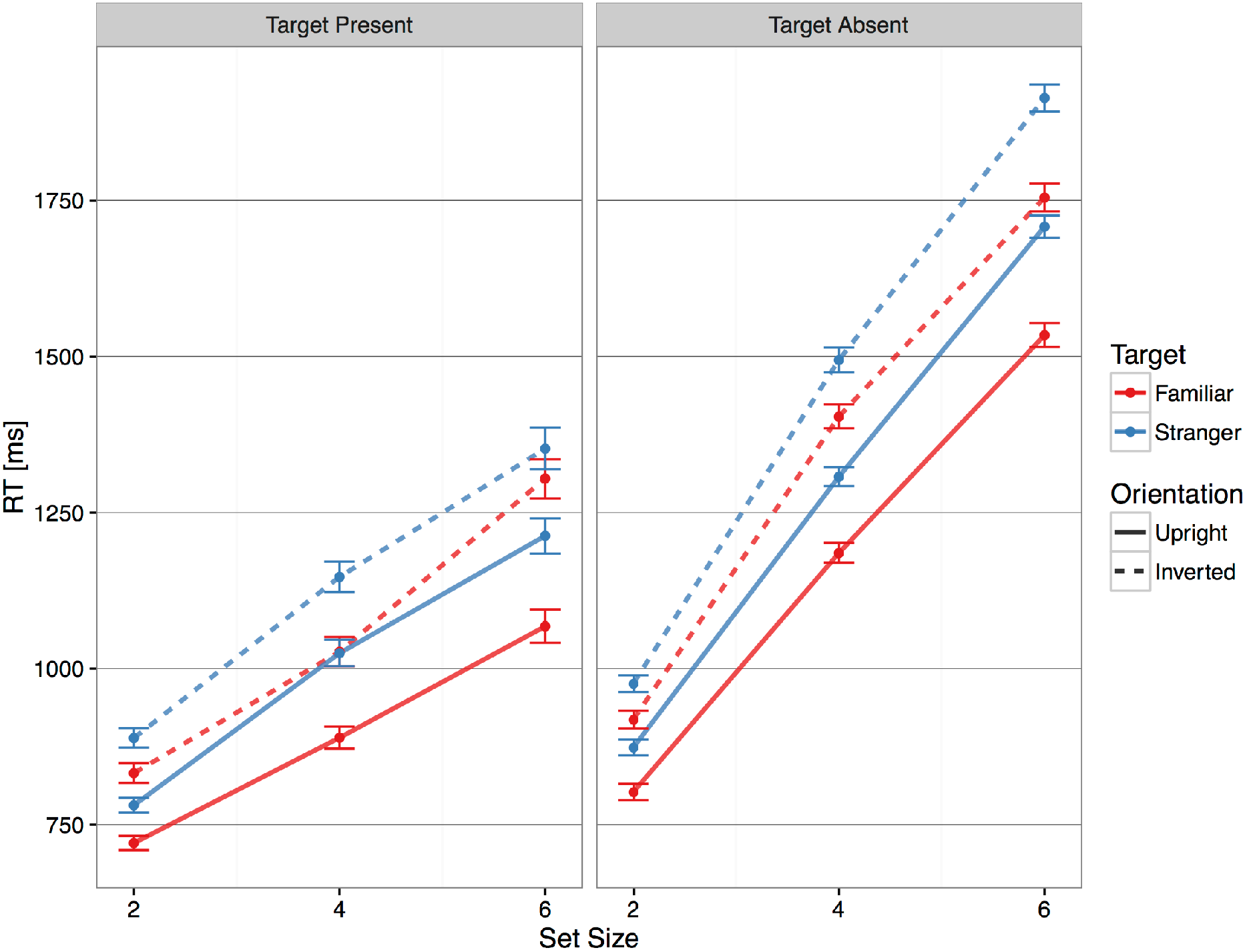
Average reaction times according to presence of the target. Subjects were always faster at determining the presence or absence of a familiar target face compared to a stranger target face. Solid lines show upright condition, dashed lines show inverted condition. Red: familiar targets; Blue: stranger targets. Error bars show 95% bootstrapped confidence intervals.

**Fig 4.**
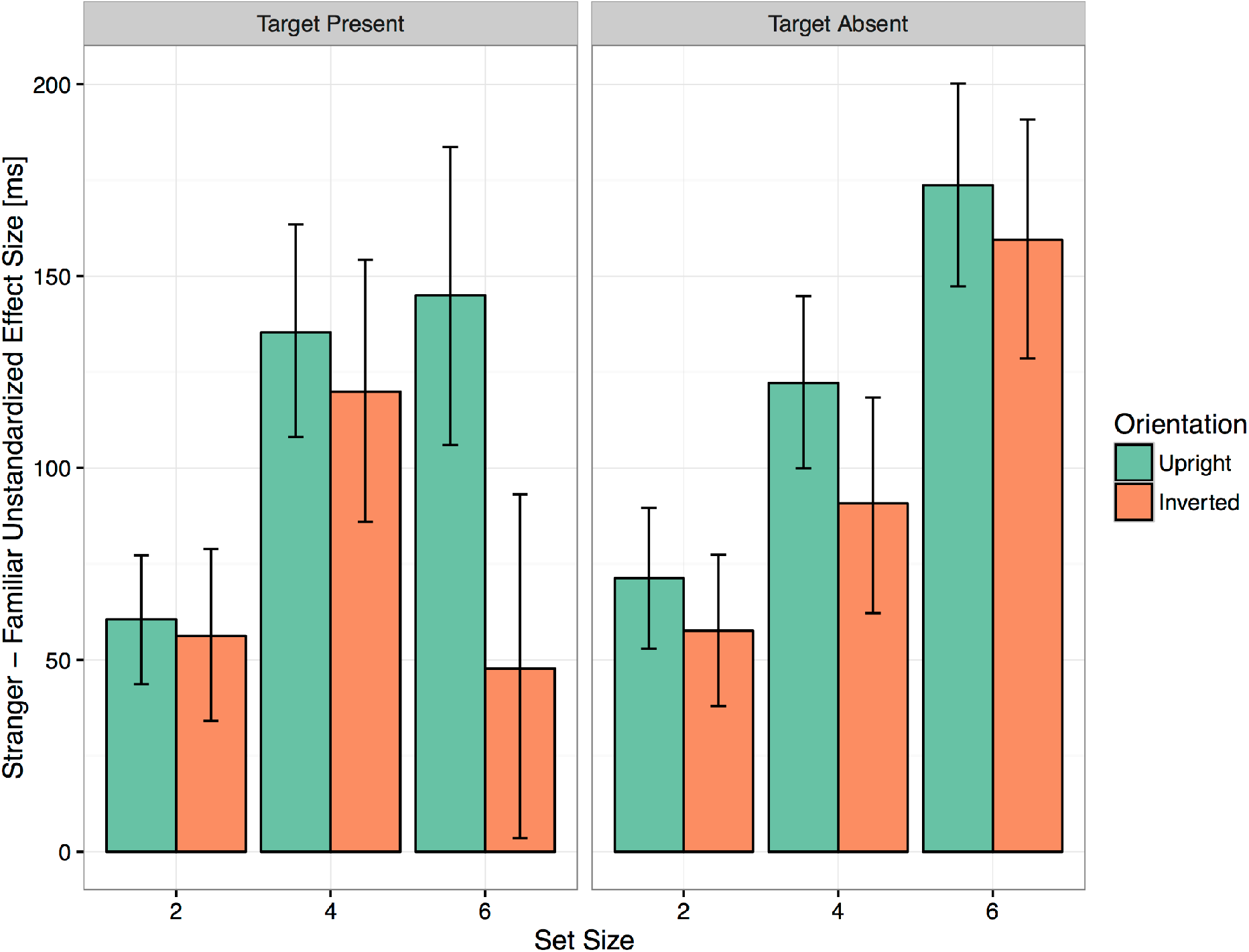
Average unstandardized effect size of Familiarity for Upright and Inverted faces in Target Present and Absent trials. Error bars show 95% bootstrapped confidence intervals.

These differences were further analyzed by looking at the estimates of Set Size 1 and slopes. With upright faces, the Set Size 1 estimates were 632 ms [615, 649] for familiar faces, and 683 ms [663, 702] for stranger faces. With inverted faces, they were 699 ms [677, 722] for familiar and 783 ms [759, 806] for stranger faces. We found a nonsignificant trend towards a greater effect of familiarity for inverted faces: 51 ms [26, 77] for upright faces, and 83 ms [50, 116] for inverted faces (difference 33 ms [-10, 74]).

The significant interaction terms in the linear mixed-effect model reflected differences in the search slopes. Search slope estimates were significantly lower for familiar faces in the upright condition: 87 ms/item [80, 94] vs. 108 ms/item [101, 116] for stranger faces (difference of 22 ms/item [11, 32]). The search slopes for inverted faces were steeper than those for upright faces, and they did not differ across familiarity (familiar faces 122 ms/item [112, 130]; stranger faces 116 ms/item [107, 125]; difference -4 ms/item [-17, 8]).

#### Target Absent Trials

All main effects of interest were significant: Set Size (*χ*^2^(2) = 8131.39, p < .001), Familiarity (*χ*^2^ (2) = 414.31, p < .001), and Target Orientation (*χ*^2^ (2) = 792.64, p < .001). The two-way interactions were significant, but the three-way interaction was not: Set Size x Familiarity (*χ*^2^(2) = 6.59, p < .05); Set Size x Target Orientation (*χ*^2^(2) = 6.20, p < .05); and Familiarity x Target Orientation (*χ*^2^(1) = 6.75 p < .01) (see S1 File). The average effect size for Familiarity was 122 ms [110, 135] in the upright condition, and 103 ms [87, 118] in the inverted condition (difference 20 ms [0, 40]). Fig 4 shows the effect size of Familiarity at each set size.

The search slopes in the target absent trials were about two times those in the target present trials, consistent with a serial self-terminating search. Interestingly, search slopes were steeper when subjects were looking for stranger targets, despite the distractors presented being the same in both familiar and stranger blocks. With upright faces, the search slope was 184 ms/item [178, 189] for familiar targets and 210 ms/item [204, 215] for stranger targets (difference 26 ms [19, 34]). The search slopes were steeper for inverted targets, but less so in familiar than stranger blocks (familiar: 210 ms/item [203, 216]; stranger: 237 ms/item [230, 243]; difference 27 ms [18, 36]).

## Discussion

In this study subjects searched for friends’ faces and strangers’ faces in a visual search task. We found a processing advantage for personally familiar faces that was robust to face inversion.

Subjects’ behavior could be framed in terms of a self-terminating serial search, with target-absent search slopes about twice the target-present ones. In target present trials subjects were highly accurate both with familiar and stranger targets, showing no evidence of a speed-accuracy trade-off.

Set Size 1 estimates showed that familiar face targets were processed faster than stranger target faces when presented both upright and inverted. This result adds to the evidence that personally familiar faces benefit from facilitated processing in a variety of experimental conditions [13–15] and real-life situations [7].

Critically, in this experiment we showed that the advantage of familiar face processing extended to inverted faces. Evidence suggests that turning a face upside-down reduces holistic perceptual processing and favors feature-based processing [18,19,24,25,29]; see also [20,21]. Thus, the faster detection of personally familiar faces in the inverted condition suggests that more efficient processing of personally familiar faces rests largely on enhanced processing of local facial features.

Our findings extend the theoretical relevance of the results by Tong and Nakayama (1999), who used subjects’ own faces as familiar identities. By using faces of subjects’ friends instead of subjects’ own faces, we made the experimental task closer to everyday experience. We spend more time looking at the faces of other people, especially personally familiar others, than at oneself, and we are more likely to search for a familiar face in a crowd rather than search for one’s own face.

We found that the search slopes differed between familiar and stranger conditions for upright faces on target present trials, and for both upright and inverted faces on target absent trials, but not for inverted faces on target present trials. These results indicate that subjects were faster at rejecting a stranger distractor when looking for a familiar face target than when looking for a stranger face target, even in target absent trials, in which the stimulus arrays were equivalent for familiar target and unfamiliar target blocks. The increase of the reaction times based on the number of items in the search array is consistent with a serial self-terminating search that was faster when searching for familiar face targets than for stranger face targets. This indicates that the internal representation of a familiar face, against which each distractor is compared, is either more robust and precise or sparser. We propose that familiarity may direct processing to specific features that are diagnostic of a familiar face’s identity, whereas the representation of a stranger’s face does not focus processing on similar diagnostic features.

Our previous results support the hypothesis for a streamlined detection of familiar faces based on diagnostic, identity-specific features. We have shown that changes in eye gaze, a local feature that serves as a potent social cue, are detected faster when conveyed by personally familiar faces [14]. We also showed that personally familiar faces are distinguished from stranger faces in a saccadic reaction time task at a latency of 180ms [15]. This very rapid detection of familiarity is faster than the time required to build a view-invariant representation of faces the monkey face patch system [48], further corroborating our hypothesis that rapid familiarity detection is based on a simpler, perhaps feature-based, process.

The slightly smaller effect of familiarity on reaction times for inverted faces than for upright faces, as reflected by the significant Familiarity x Orientation interaction, may suggest that some of the features of familiar face representations that afford more rapid processing are configural or holistic. However, the greater magnitude of the familiar advantage even for the inverted faces shows that this facilitation relies mainly on local features. Related work by others also indicates that configural information is less important for recognition of a familiar identity (see [12] for a cogent argument).

In summary, the results of our experiment add to the existing evidence that the human visual system is finely tuned for rapid detection and identification of familiar faces, much more so than of stranger faces. Participants searched for a familiar or stranger identity among distractors presented in either an upright or inverted orientation. They responded faster when searching for familiar faces even in the inverted condition. Taken together, our results suggest that robust representations for familiar faces contain information about idiosyncratic facial features that allow subjects to detect or reject identities when searching for a friend’s face in a crowd of stranger faces.

## Acknowledgements

We would like to thank Sebastian M. Frank, J. Swaroop Guntupalli, the members of the Gobbini lab and Haxby lab for helpful discussions on these results.

